# Maternal transmission of a paramutant phenotype requires intact DNMT2 functions in the male germline

**DOI:** 10.1101/2020.10.22.351155

**Authors:** Tian Yu, Yeming Xie, Chong Tang, Yue Wang, Shuiqiao Yuan, Huili Zheng, Wei Yan

## Abstract

The molecular mechanism underlying epigenetic inheritance remains largely unknown. Previous studies have shown that transmission of paternal acquired traits through the male germline (i.e., sperm) requires functional DNMT2 to maintain normal profiles of sperm-borne tRNA-derived small RNAs (tsRNAs). Here we report that maternal transmission of a *Kit* paramutant phenotype (white tail tip) through the female germline (i.e., oocytes) also requires normal function of DNMT2 and normal profiles of DNMT2-dependent tsRNAs and other small noncoding RNAs (sncRNAs) in sperm. Specifically, ablation of DNMT2 leads to aberrant profiles of tsRNAs and other sncRNAs in sperm, which correlate with drastically dysregulated mRNA transcriptome in pronuclear zygotes derived from oocytes carrying the *Kit* paramutation and a complete blockage of transmission of the paramutant phenotype through the oocytes. Together with previous reports, the present study suggests that both paternal and maternal transmission of epigenetic phenotypes requires DNMT2-dependent tsRNAs in sperm.

## INTRODUCTION

DNA methylation, as a major epigenetic modification in the genome of higher eukaryotes, is catalyzed by DNA methyltransferase enzymes (DNMTs) by addition of a methyl group to the 5th carbon position of cytosine, resulting 5-methylcytosine (5mC) (*1, 2*). DNMT1 is the most abundant DNA methyltransferase in mammalian cells, and mainly methylate hemi-methylated CpG dinucleotides as a maintenance methyltransferase, although DNMT1 has also been shown to function as a *de novo* DNA methyltransferases (*3*). Both DNMT3a and DNMT3b are able to methylate previous unmethylated CpG sequences and thus, are called *de novo* methyltransferases (*1, 2*). In addition to DNMT1, DNMT3a and DNMRT3b, there are two noncanonical family members, DNMTand DNMT3L, neither of which has DNA methyltransferase activity (*4*). DNMT3L appears to interact with DNMT3a and DNMT3b to regulate their DNA methyltransferases activities (*5*), whereas DNMT2 has been found to act as a tRNA methyltransferase, responsible for catalyzing methylation of cytosine 38 in the anticodon loop of tRNA^Asp^_GUC_ (*6*). DNMT2-dependent methylation protects the anticodon loop of tRNAs from cleavage in flies and mice, and a lack of m5C at position 38 of three tRNAs (tRNAAsp_GUC_, tRNAGly_GCC_ and tRNAVal_AAC_) induces fragmentation of these tRNAs, leading to production of tRNA fragments (tsRNAs), also called tRNA-derived small RNAs (tsRNAs) (*7, 8*). It has been reported that tsRNAs are highly abundant in sperm (*9*). More interestingly, the tsRNA profiles are altered in sperm of the male mice with high fat diet-induced metabolic disorders, and injection of small RNA fractions containing tsRNAs purified from these male mice into the wild-type zygotes recapitulates the metabolic disorder phenotype in offspring (*10*). A related study also reports that the sperm may have gained a large number of the tsRNAs from the epididymal exosomes (called epididymosomes) secreted by the epididymal epithelial cells during sperm transit through the epididymis, and sperm tsRNA profiles get alerted in sperm from male mice on a protein-restricted diet (Sharma et al., 2016). These data strongly suggest that sperm-borne tsRNAs may function as a messenger to transmit information on specific epigenetic alterations/epimutations induced by various factors, e.g., environmental exposure, drug, infection, lifestyle changes etc., to offspring (*11*). Therefore, it is critical to understand how tsRNAs are produced and how these tRNA fragments act at molecular levels to convey epigenetic information during fertilization and early embryonic development.

Given that DNMT2 is involved in tsRNA biogenesis by generating 5mC on C38 of three substrate tRNAs (*6–8*), we and others have explored the impact of *Dnmt2* ablation on epigenetic inheritance. The data indicate that *Dnmt2* inactivation in male mice can block, to a large extend, the paternal/sperm-mediated transmission of acquired phenotypes, including the while tails in *Kit* paramutant mice and HFD-induced metabolic disorders (*12, 13*). Therefore, tsRNAs appear to be required for efficient transmission of acquired paternal phenotypes through sperm. However, many questions remain, e.g., how does ablation of DNMT2 affect sperm-borne tsRNA contents? What is the impact of DNMT2 ablation on the profiles of other sperm-borne sncRNAs? Are normal sperm-borne tsRNAs required for efficient transmission of a maternal epigenetic phenotype through oocytes/eggs? How do sperm-borne tsRNAs and other sncRNAs act to regulate gene expression during zygotic genome activation in pronuclear embryos? The present study aimed to address these fundamental questions.

We adopted a *Kit* paramutation mouse model because the phenotype induced by the paramutant wild-type allele, i.e., white tail tip (WTT), is easy to see and measure (*14*). Using WTT as a phenotypic readout, we set out and tested the effects of *Dnmt2*-null sperm on the maternal transmission efficiency of the WTT phenotype. To our surprise, our breeding data revealed that DNMT2 ablation in sperm almost completely abolished maternal transmission of the WTT phenotype. Further analyses suggest that sperm-borne, DNMT2-dependent tsRNAs and other small RNAs are essential for the transmission of the *Kit* paramutant phenotype through the maternal germline (i.e., oocytes). Comprehensive examination of small RNA methylome and large and small RNA transcriptomes of DNMT2-*null* sperm and 3PN and 4PN zygotes derived from wildtype or *Kit^+/copGFP^* oocytes injected with *Dnmt2*-null or wildtype sperm led us to conclude that DNMT2-depdent tsRNAs and miRNAs regulates genes involved in histone modifications and chromatin organization during early embryonic development and are required for proper transmission of maternal epimutations.

## Results

### Inhibition of maternal transmission of the *Kit* paramutant phenotype by *Dnmt2*-null sperm

Insertion of a reporter gene cassette, either *LacZ* (*15*) or *copGFP* (*14*), into exon1 of *Kit* has been shown to induce coat color changes in heterozygous mice of C57Bl/6J strain, characterized by white patches in the abdomen and the tail (Fig. 1a). Interestingly, wildtype (WT) offspring derived from heterozygous (*Kit^+/copGFP^*) parents tend to display white tail tips (WTT) that are highly variable in length (Fig. 1b). The white tail tips represent a specific phenotype induced by the paramutant *Kit* allele, i.e., the WT allele derived from the *Kit^+/copGFP^* parent (*14, 15*). The WTT phenotype can be transmitted *via* both paternal and maternal germlines and maintained across multiple generations (*14*). In our WT C57Bl6/J mouse breeding colonies (totally unrelated to the *Kit*^*copGFP*^ line), ~30% of the mice displayed WTTs, including 29% showing shorter and only 1% with longer WTTs (Fig. 1c), which serve as baseline penetrance (~30%) and severity (< 3mm) of the WTT phenotype among WT C57Bl/6J mice in our lab. When WT male mice with black tail tips (BTT) were bred with *Kit^+/copGFP^* female mice, ~60% of the WT offspring had WTTs, including 31% showing longer (>3mm) and 28% with shorter WTTs (Fig. 1c, left upper panel). These results suggest that the WT paramutant *Kit* allele (the maternal WT *Kit* allele bearing epimutations) can be transmitted to offspring through the maternal germline (oocytes), and the WTT phenotype induced by the *Kit* paramutant allele is often more severe (>3mm in length). Intriguingly, when the WT BTT males were switched to *Dnmt2*-null BTT males (*Dnmt2*^*-/-*^ males x *Kit^+/copGFP^* females), the maternal transmission of the paramutation-induced WTT phenotype was largely abolished, as evidenced by the significantly reduced penetrance of the WTT phenotype (59% → 35%) and decreased severity (31% → 3%) in *Kit* wildtype offspring (Fig. 1c, right lower panel). Interestingly, both the phenotypic penetrance and severity were similar to those in offspring of WT BTT breeding pairs (Fig. 1 c, left upper panel). Furthermore, when WT BTT females were bred with *Dnmt2*-null BTT males, the penetrance of the WTT phenotype and severity in the *Dnmt2^+/-^* offspring with WT *Kit* locus appeared to be reduced slightly compared to the pure WT breeding scheme (WT BTT x WT BTT) (Fig. 1c, right upper panel). The common finding lies in that the WTT phenotype appears to be inhibited when the males are on the *Dnmt2*-null background. Although the F1s studied were all wild-type in *Kit* locus, those fathered by *Dnmt2*-null males are heterozygous in *Dnmt2* locus (*Dnmt2^+/-^*), raising the possibility that the reduced penetrance of the WTT phenotype may be due to the *Dnmt2* heterozygosity (*Dnmt2^+/-^*), rather than a lack of DNMT2-dependent tsRNAs and/or other sncRNAs in *Dnmt2*-null sperm. In other words, one allele of *Dnmt2* in the WT *Kit* early embryos may not be sufficient to support efficient transmission of the WTT phenotype carried by the maternal WT *Kit* allele. To test this possibility, we set up breeding pairs using *Dnmt2^+/-^* males and *Kit+/copGFP* females to examine the penetrance of the WTT phenotype in offspring that are WT in both *Kit* and *Dnmt2* loci, as well as offspring that are WT in *Kit* locus, but heterozygous in *Dnmt2* locus. Interestingly, the distribution of the phenotypic penetrance in both *Dnmt2^+/+^; Kit^+/+^* and *Dnmt2^+/-^; Kit^+/+^* F1s was similar to that in F1s derived from WT males and *Kit^+/copGFP^* females, all with increased proportions of the longer WTT (Supplemental information, Fig. S1). This finding strongly suggests that *Dnmt2* heterozygosity in early WT *Kit* embryos does not affect the efficiency of maternal transmission of the WTT phenotype carried on the maternal WT *Kit* allele because both *Dnmt2^+/+^* and *Dnmt2^+/-^* F1s displayed the same enhanced penetrance of the longer WTT phenotype. Thus, it is highly likely that *Dnmt2*-null sperm, rather than *Dnmt2* heterozygosity of F1 embryos, are responsible for the inhibitory effects on maternal transmission of the WTT phenotype.

**Fig.1.**
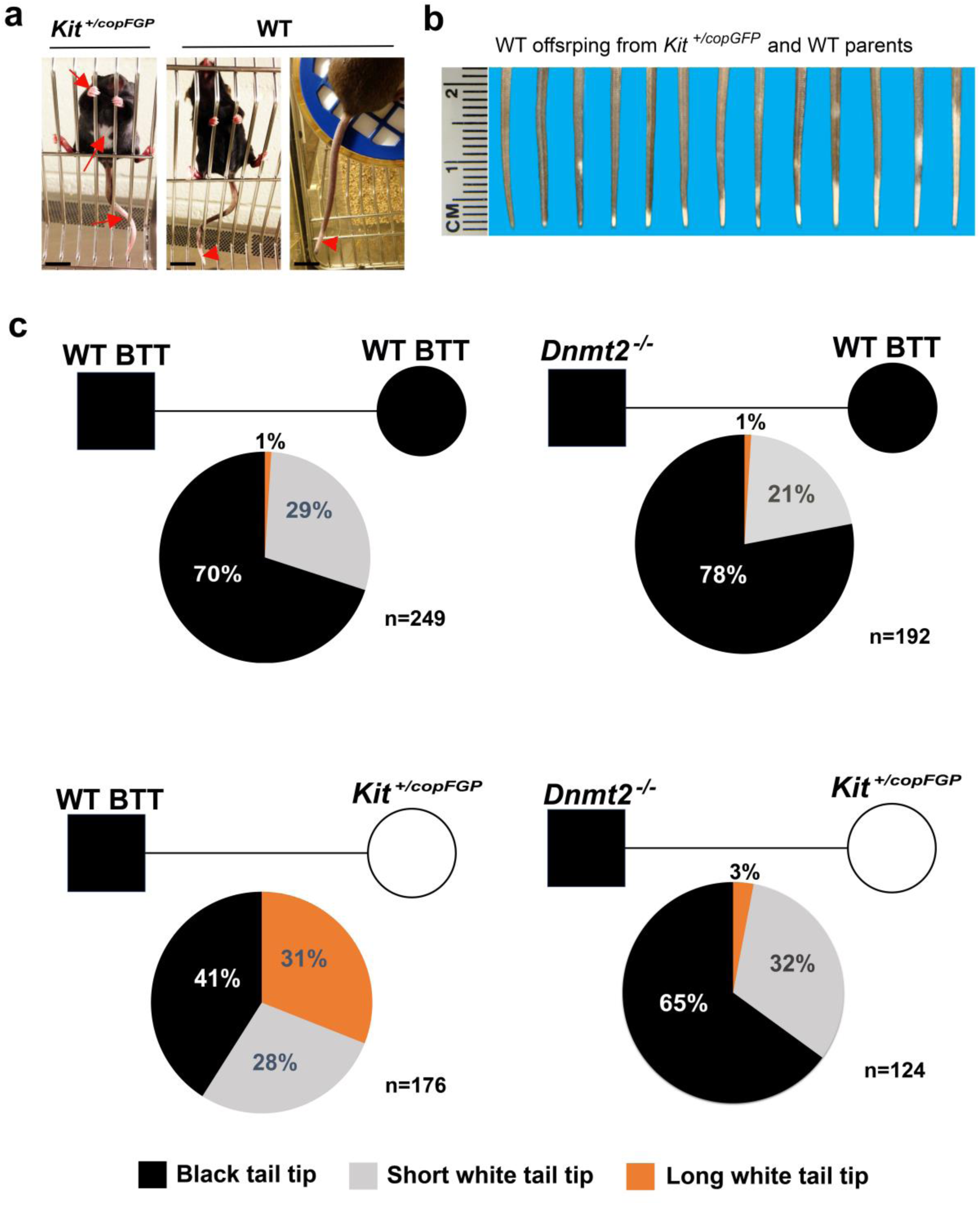
Blockage of maternal transmission of a Kit paramutant phenotype by *Dnmt2*-null sperm. (a) The *Kit^+/copGFP^* mice display white paws, long white tail tips and white abdominal patches (arrows), whereas wild-type (WT) offspring derived from either *Kit^+/copGFP^* x *Kit^+/copGFP^* or WT x *Kit^+/copGFP^* breeding pairs tend to have white tail tips (WTT) (arrowheads). In this *Kit* paramutation mouse model, the wild-type *Kit* allele in the *Kit^+/copGFP^* mice has been epigenetically modified such that inheritance of this paramutant wild-type Kit allele can induce the WTT phenotype in wild-type offspring. Scale bar = 5 mm. (b) Variable severity of the WTT phenotype in WT offspring of *Kit^+/copGFP^* x *Kit^+/copGFP^* or WT x *Kit^+/copGFP^* parents. In the present study, long and short white tail tips were defined by the cumulative length of WTT >0.5 cm and <0.5 cm, respectively. (c) Pie charts showing the penetrance of the WTT phenotype in *Kit* WT offspring derived from four breeding schemes. The number (n) of mice observed are marked.

### Altered tsRNA profiles characterized by decreased m5C levels at C38, increased tsRNA levels and global shortening of tsRNAs in *Dnmt2*-null sperm

Unlike DNMT1 and DNMT3, DNMT2 catalyzes the 5’ cytosine methylation (m5C) at position 38 (C38) of three major tRNAs: Val_AAC_, Asp_GUC_, and Gly_GCC_ (*4*) (Fig. 2a). Proper C38 methylation protects these tRNAs from degradation into tsRNAs (*7, 8*). Thus, DNMT2 is believed to be a critical regulator of tsRNA biogenesis (*4, 6*). To determine the changes in tsRNA profiles in *Dnmt2*-null sperm, we performed high-throughput small RNA bisulfite sequencing (sRNA-BS-Seq). The cytosine on tRNA reads were identified first and the percentage of m5C (methylated counts/total read counts for each tRNA) was calculated. While the C38 in all 7 isoforms of the three DNMT2 substrates (Val_AAC_, Asp_GUC_, and Gly_GCC_) was hypermethylated in WT sperm, it was hypomethylated in *Dnmt2*-null sperm (Fig. 2b, the left panel). It has been shown that lack of C38 5mC, tRNAs tend to degrade (*4, 6*). Indeed, we observed significant reduction in the overall levels of all the 7 DNMT2 substrate tRNAs in *Dnmt2*-null sperm compared to WT sperm (Fig. 2b, the middle panel). Consistent with previous data (*4*), as a non-DNMT2 substrate, tRNA Glu_UUC_ showed no significant changes in either methylation levels at C38 or total read counts in *Dnmt2*-null sperm (Fig. 2b, the middle panel), validating our quantitative analyses. While tRNA degradation appeared to be enhanced, levels of tsRNAs derived from DNMT2 substrate tRNAs were increased in *Dnmt2*-null sperm (Fig. 2b, the right panel). Moreover, the average length of tsRNAs derived from the three major DNMT2 substrate tRNAs appeared to be significantly reduced in *Dnmt2*-null sperm compared to WT sperm (Fig. 2c and Supplemental information, Fig. S2). Although the length reduction was the most prominent in tsRNAs derived from the DNMT2 substrate tRNAs, those from non-DNMT2 substrate tRNAs, e.g., Glu_UUC_, also displayed a minor length reduction despite comparable abundance (Fig. 2c). Indeed, the shortening in tsRNAs appeared to be global in *Dnmt2*-null sperm (Fig. 2d). The global tsRNA shortening suggests that tRNAs may not break at the canonical site (i.e., C38) in *Dnmt2*-null sperm. To test this notion, we analyzed the abundance of individual nucleotides by plotting the normalized percentage abundance of each nucleotide against each position along the full-length tRNAs (Fig. 2e). Remarkably, a stretch of nucleotides around the anticodon region (27-45nt) of tRNAs were drastically underrepresented (arrow in Fig. 2e), suggesting that the decreased m5C on C38 may have destabilized its surrounding anticodon region, leading to enhanced degradation, which may have contributed to the global shortening of tsRNAs in *Dnmt2*-null sperm. Moreover, 5’ tsRNAs appeared to be more abundant than 3’ tsRNAs in WT sperm, and this bias became more prominent in *Dnmt2*-null sperm (Fig. 2e). To corroborate this finding, we analyzed the small RNA-Seq data from another publication (Yang et al., 2016), and obtained similar results showing 5’ tsRNAs were dominant in both sperm and zygotes in mice (Supplemental information, Fig. S3). These observations are consistent with Northern blot analyses showing that 5’ tsRNAs are much more abundant than 3’ tsRNAs (*16*). Together, out data support the role of DNMT2 as a methyltransferase responsible for the C38 5mC of tRNAs, and a lack of this mark leads to enhanced breakups of not only the three substrate tRNAs at C38, but also the other tRNAs around the anticodon loop region (between positions 27-45nt).

**Fig.2.**
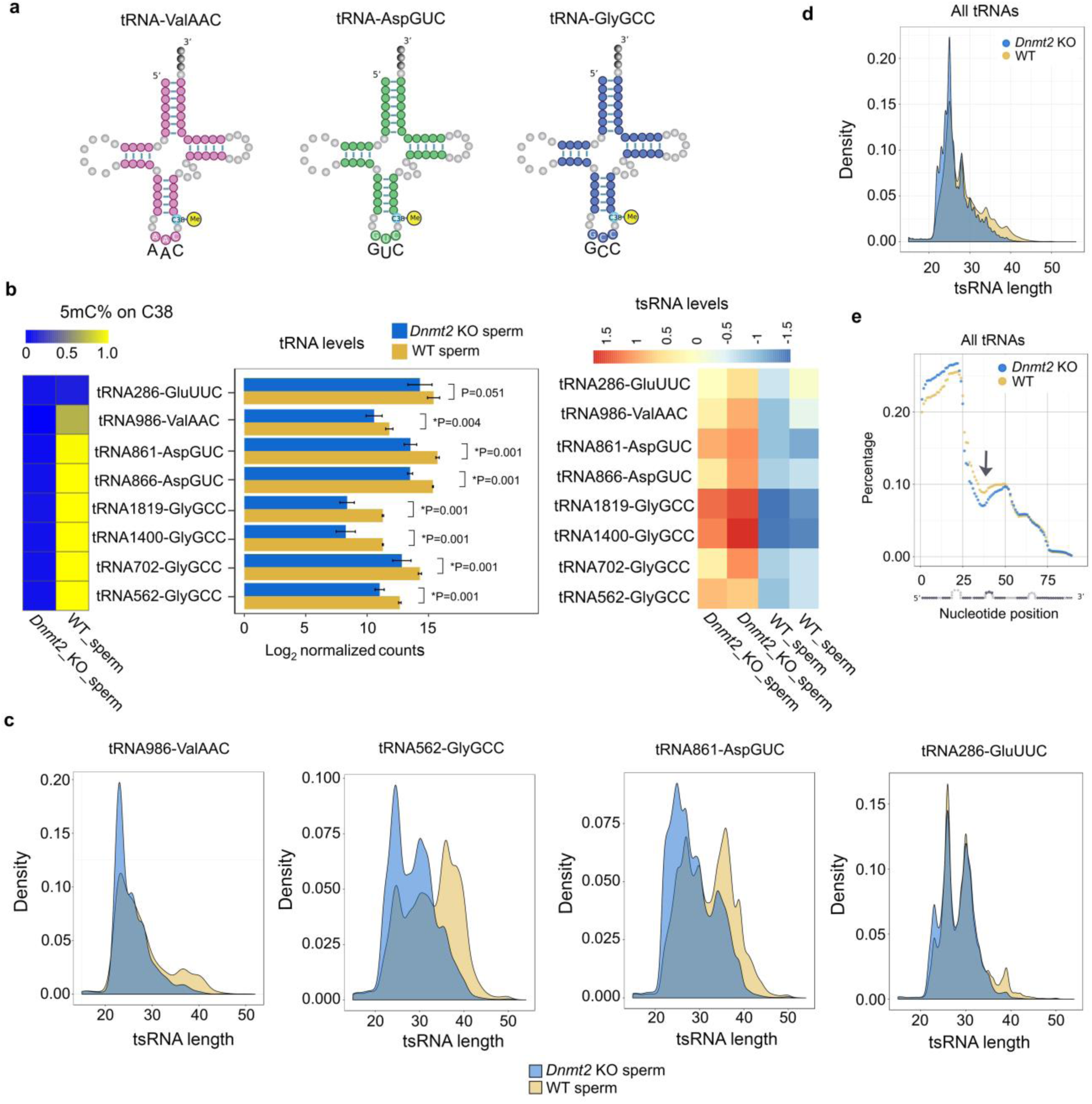
Altered tsRNA profiles in *Dnmt2*-null sperm characterized by decreased m5C levels on C38 of tRNAs, reduced substrate tRNA levels, increased tsRNA levels and global shortening of tsRNAs. (a) Schematics of the 3 confirmed substrate tRNAs of DNMT2 (tRNA-Val_AAC_, Asp_GUC_ and Gly_GCC_) in mice. (b) Levels of 5mC, substrate tRNAs (seven isoforms) and tsRNAs (derived from the three substrate tRNAs) in *Dnmt2* KO and wildtype (WT) sperm. Heatmap (left panels) shows the mean percentage of m5C on C38 of tRNAs, and the bar graphs (middle panels) indicate the log2 values of normalized counts to represent tRNA levels in *Dnmt2* KO and WT sperm. The relative levels of substrate tRNAs-derived tsRNAs represent normalized tsRNA counts/ normalized tRNA counts (heatmap in the right panels). Each row represents a subtype from the 3 DNMT2 substrate tRNAs and one non-DNMT2 substrate tRNA (tRNA286-Glu_UUC_). The data were presented as means ± SD from biological replicates, *n*=2. Sums of two technical repeats in each of the biological replicates were used for analyses. *: p<0.05. (c) Density plots showing the length distribution of tsRNAs derived from three major DNMT2 substrate tRNAs (tRNA986-Val_AAC_, tRNA562-Gly_GCC_, and tRNA861-Asp_GUC_) and one non-DNMT2 substrate (tRNA286-Glu_UUC_) in *Dnmt2* KO and WT sperm. (d) Density plots showing the length distribution of all of the tsRNAs detected by sncRNA-BS-seq. (e) Distribution of individual nucleotides along the entire length of tRNAs. The normalized percentage abundance (y axis) was plotted against positions of each nucleotide (x axis). Note that a stretch of nucleotides around the anticodon region (27-45nt) of tRNAs was drastically underrepresented (arrow), and that 5’ tsRNAs appeared to be more abundant than 3’tsRNAs in WT sperm, and this bias became more evident in *Dnmt2*-null sperm.

### Expression profiles of other small noncoding RNAs are also altered in *Dnmt2*-null sperm

To explore the secondary effect of *Dnmt2* inactivation on sncRNA expression profiles, we annotated eight known sncRNAs, including endo-siRNAs, miRNAs, mitochondrial DNA-encoded rRNA (mt rRNA) and tRNAs (mt tRNA), piRNAs, rRNAs-derived small RNAs (rsRNAs), snoRNAs, and snRNAs using AASRA (*17*), and compared their normalized total counts between *Dnmt2*-null and WT sperm. Differential expression was observed in all 8 sncRNA species between *Dnmt2*-null and WT sperm (Fig 3a). For example, levels of tRNAs were lower in *Dnmt2*-null than in WT sperm, likely due to the enhanced tRNA breakups into tsRNAs in the absence of DNMT2. In contrast, miRNA levels were upregulated in *Dnmt2*-null sperm. Remarkably, among all the dysregulated sncRNAs, miRNAs were mostly upregulated, while levels of intact tRNAs were mainly downregulated in *Dnmt2*-null sperm, as compared to WT sperm (Fig 3b). Indeed, 26 out of 30 miRNAs that were differentially expressed between *Dnmt2*-null and WT sperm were up-regulated (Fig 3c), whereas the total tRNAs with significant fold changes were down-regulated (Fig. 3d and Supplemental information, Table S1) in *Dnmt2*-null sperm. Accordingly, levels of both substrate and non-substrate tsRNAs were all upregulated in *Dnmt2*-null sperm, as compared to WT sperm (Fig. 3d). Expression profiles of other sncRNAs, including piRNAs, endo-siRNAs, snoRNAs, snRNAs and rsRNAs, were also altered (Supplemental information, Fig S4 and Table S2).

Given that these differentially expressed sncRNAs all resulted from *Dnmt2* inactivation and all may participate in the regulation of gene expression during and post fertilization, we termed them “DNMT2-dependent sncRNAs”. By comparing the DNMT2-dependent sncRNAs with those expressed in oocytes (*18*), we found that ~23% of the DNMT2-dependent sperm miRNAs and 55% of the DNMT2-dependent sperm tRNAs were shared with the oocytes (Fig. 3f). Among the shared DNMT2-dependent tRNAs, the majority (~79%) displayed greater normalized counts in sperm (Fig. 3g and Supplemental information, Table S3). Three out of the top 7 DNMT2-regulated miRNAs showed higher expression levels in sperm than in oocytes (Fig. 3h). Together, these data suggest that the majority of the DNMT2-dependent small RNAs are enriched in sperm. To validate the dysregulation of miRNAs, we analyzed levels of 8 DNMT2-dependent miRNAs (miR-34b-3p, 701-3p, 96-5p, 5108, 8104, 5106, 6240, and 6958-5p) in WT, *Dnmt2^+/-^* and *Dnmt2*^*-/-*^ sperm using qPCR, and found that levels of these 8 miRNAs were similar in both WT and *Dnmt2^+/-^* sperm, but much higher in *Dnmt2*-null sperm (Supplemental information, Fig. S5). The data further support the notion that it’s the sperm-borne, DNMT2-dependent tsRNAs and other sncRNAs that block maternal transmission of the WTT phenotype.

### Transcriptomic alterations upon zygotic genome activation in PN3/PN4 prenuclear embryos derived from *Kit*^*+/copGFP*^ oocytes injected with either *Dnmt2*-null or WT sperm

. The major difference between the two breeding schemes (i.e., *Dnmt2*-null males X *Kit^+/copGFP^* females vs. WT males X *Kit^+/copGFP^* females) lies in the sperm (*Dnmt2*-null vs. WT). Therefore, the cause for failed maternal transmission of the paramutant WTT phenotype must be hidden within the *Dnmt2*-null sperm. Given that DNMT2 methylates tRNAs and the profiles of tsRNAs and other sncRNA species are altered in *Dnmt2*-null sperm, it is plausible to hypothesize that the *Dnmt2*-null sperm, due to their altered sncRNA profiles, exert effects on the transcriptome of early embryos and thus, block the maternal transmission of the WTT phenotype. To test this hypothesis, we injected WT or *Dnmt2*-null sperm into the oocytes from either *Kit^+/copGFP^* or WT mice, and collected zygotes at PN3 and PN4 stages for single cell RNA-seq analyses (Fig. 4a). We chose to analyze the transcriptome of zygotes at pronuclear PN3/PN4 stages because the paternal genome has been activated with greater transcriptional activities than the maternal genome at these stages (*19*). When WT and *Dnmt2*-null sperm were injected into *Kit^+/copGFP^* oocytes, 50% of the resulting embryos, in theory, would be heterozygous for *Kit* (*Kit^+/copGFP^*), which were excluded based on microscopic observation of copGFP expression and PCR detection of copGFP transcripts using total cDNAs prepared from single zygotes. Only *Kit^+/+^* zygotes were selected for single cell RNA-seq analyses.

**Fig. 3.**
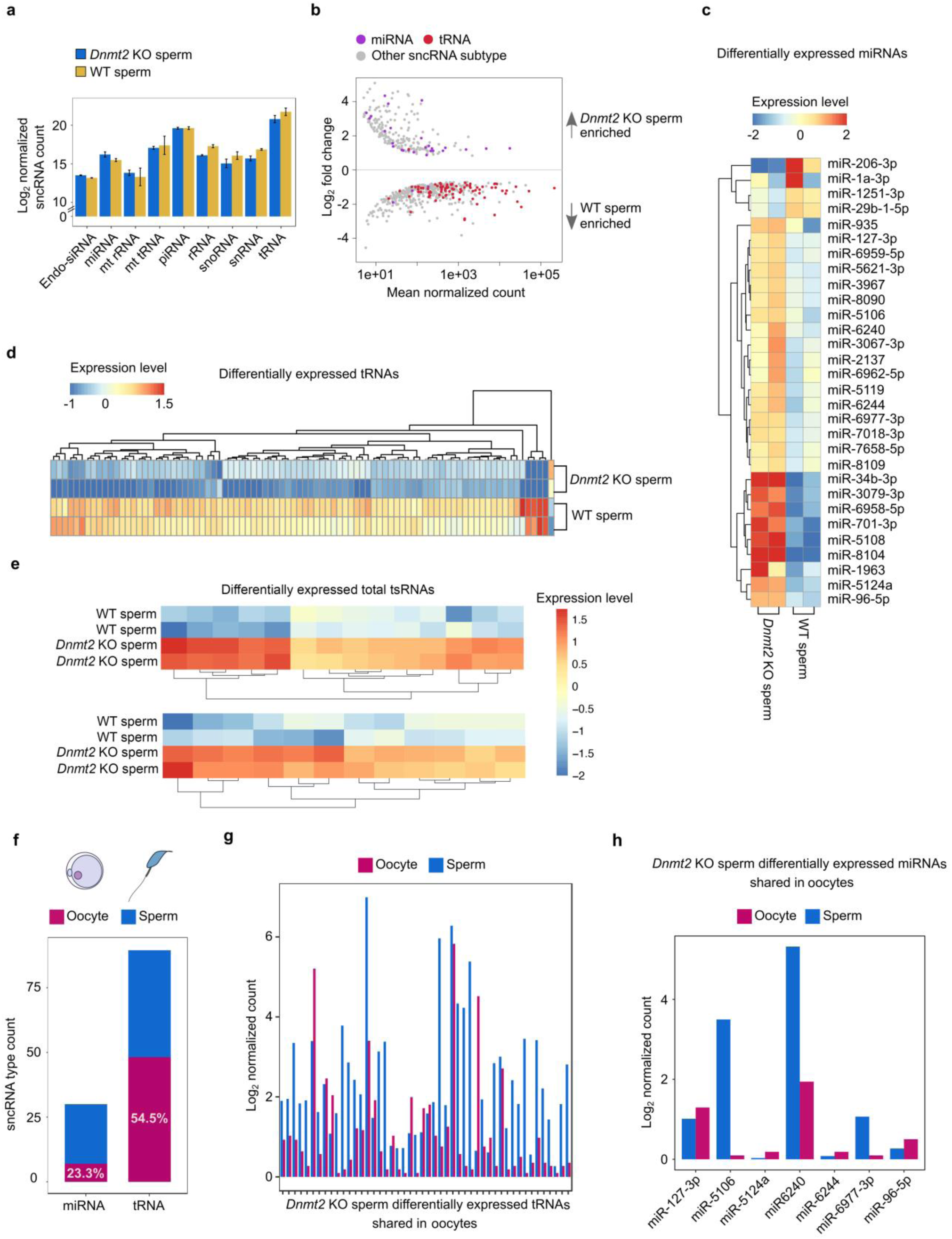
Altered profiles of other sperm-borne sncRNAs in *Dnmt2*-null sperm. (a) Total levels of 9 types of sncRNAs in *Dnmt2* KO and WT sperm. sncRNA reads were identified from sncRNA-Seq data in two biological replicates with two technical repeats, followed by DESeq2 log2 normalization to generate the total counts of each sncRNA subtype. (b) MA plots showing differentially expressed (p<0.05) miRNAs, tRNAs and other sncRNAs between WT and *Dnmt2* KO sperm. Mean normalized counts represent the mean value of DESeq2 normalized counts (>5) between WT and *Dnmt2* KO sperm. Log2 fold change was calculated by the Log2 sncRNA counts of *Dnmt2* KO sperm/WT sperm. sncRNAs-Seq data were from two biological replicates with two technical repeats (n=2). (c) Heatmap showing differentially expressed miRNAs between WT and *Dnmt2* KO sperm (p< 0.05). The relative expression levels are presented by the mean-centered and normalized log-expression values. (d) Heatmap showing differentially expressed tsRNA subtypes between WT and *Dnmt2* KO sperm (p< 0.05). The relative expression levels are presented by the mean-centered and normalized log-expression values. (e) Bar graphs showing the percentage of miRNAs and tsRNA subtypes that are differentially expressed (p< 0.05) in *Dnmt2*-null sperm, but also present in oocytes. (f)) Bar graphs showing the relative expression levels of tsRNA subtypes that are not only differentially expressed (p<0.05) in *Dnmt2*-null sperm, but also present in oocytes. The *x-axis* indicates the individual tsRNA subtypes, whereas the *y-axis* shows Log2 of DESeq2-normalized read counts. (g) Bar graphs showing the relative expression levels of miRNAs that are not only differentially expressed (p< 0.05) in *Dnmt2*-null sperm, but also present in oocytes. The *x-axis* indicates the individual miRNAs, whereas the *y-axis* shows Log2 of DESeq2-normalized read counts.

**Fig. 4.**
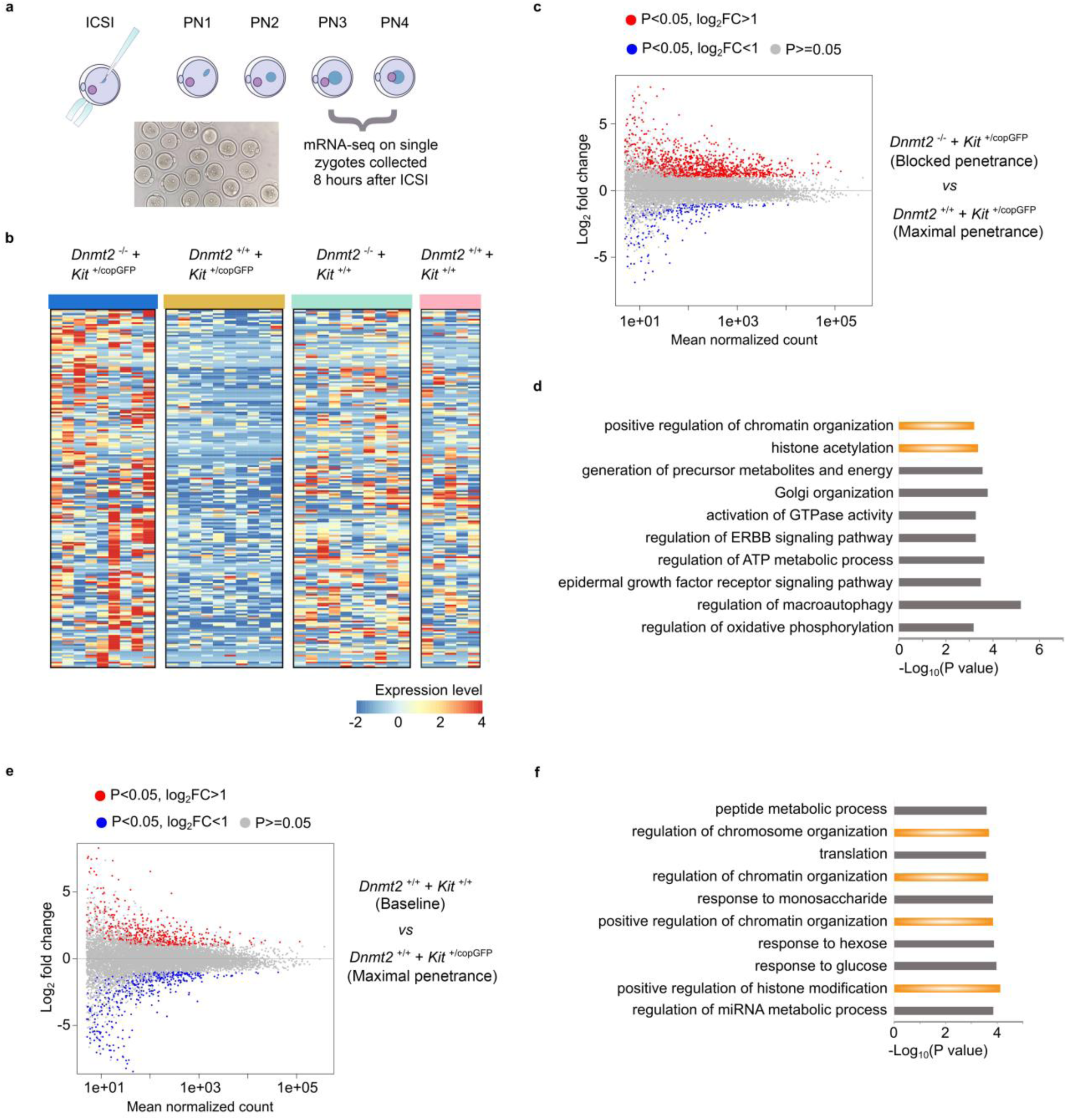
Dynamic transcriptomic changes in the pronuclear zygotes derived from *Kit*^*+/copGFP*^ oocytes and *Dnmt2*-null sperm. (a) Schematics showing the workflow of transcriptomic analyses on single zygotes of pronuclear stages 3 and 4 (PN3/PN4) produced by intracytoplasmic sperm injection (ICSI). (b) Heatmap showing the top 200 differentially expressed genes (DEGs) (p<0.05, Log_2__fold change) between *Dnmt2*^*+/-*^ zygotes from *Kit^+/copGFP^* oocytes injected with *Dnmt2*-null sperm and WT zygotes from *Kit^+/copGFP^* oocytes injected with WT sperm (Left two panels). The expression levels of these 200 DEGs in *Dnmt2^+/-^* zygotes from WT oocytes injected with *Dnmt2*-null and WT zygotes from WT oocytes injected with WT sperm (right two panels). The four types of zygotes mirror the four types of F1 mice used to analyze the penetrance of the WTT phenotype, as shown in Fig. 1C. (c) MA plots showing the DEGs in zygotes derived from *Kit^+/copGFP^* oocytes injected with *Dnmt2*-null (n=9) *or* WT (n=10) sperm. (d) Gene ontology (GO) analyses on genes dysregulated in zygotes from *Kit^+/copGFP^* oocytes injected with *Dnmt2*-null sperm compared to those from *Kit^+/copGFP^* oocytes injected with WT sperm. Top 10 enriched gene ontology terms are shown in bar graphs with −Log10 P values indicated. (e) MA plots showing the DEGs in zygotes derived from *Kit^>+/copGFP^* (n=9) or WT (n=5) oocytes injected with WT sperm. (f) Gene ontology (GO) analyses on genes dysregulated in zygotes derived from WT oocytes injected with WT sperm compared to those from *Kit^+/copGFP^* oocytes injected with WT sperm. Top 10 enriched gene ontology terms are shown in bar graphs with −Log_10_ P values indicated.

A total of four types of single PN3/PN4 stage zygotes were used for RNA-seq: *Dnmt2^+/-^*; *Kit^+/+^* PN3/4 embryos from *Kit^+/copGFP^* oocytes injected with *Dnmt2*-null sperm, *Dnmt2+/+*; *Kit^+/+^* PN3/4 embryos from *Kit^+/copGFP^* oocytes injected with WT sperm, *Dnmt2^+/-^*; *Kit^+/+^* PN3/4 embryos from WT oocytes injected with WT sperm, and *Dnmt2+/+*; *Kit^+/+^* PN3/4 embryos from WT oocytes injected with WT sperm. The four types of pronuclear embryos mirror the F1 offspring from the four breeding schemes used (Fig. 1c). A total of eighty single zygotes (20 for each group) were sequenced, and 34 that passed the quality control were used for bioinformatic analyses to identify differentially expressed genes (DEGs) (Supplemental information, Fig. S6). By calibrating against the median expression values among the four types of single zygotes, we chose the top 200 dysregulated genes (p<0.05, Log2_fold change) between zygotes from the two mating groups (*Dnmt2^+/-^*; *Kit^+/+^* PN3/4 embryos from *Kit^+/copGFP^* oocytes injected with *Dnmt2*-null sperm, and WT PN3/4 embryos from *Kit^+/copGFP^* oocytes injected with WT sperm), and created heatmaps to show the DEG patterns (Fig. 4b, left two panels, and supplemental information, Table S6). As a comparison, we also generated the heatmaps for the other two groups of zygotes (*Dnmt2^+/-^*; *Kit^+/+^* PN3/4 embryos from WT oocytes injected with WT sperm, and WT PN3/4 embryos from WT oocytes injected with WT sperm) (Fig. 4b, right two panels). In general, similar DEG profiles were observed between zygotes derived from WT oocytes injected with either WT or *Dnmt2-null* sperm (Fig. 4b, right two panels), whereas the DGE profiles were drastically different between zygotes from the *Kit^+/copGFP^* oocytes injected with WT and *Dnmt2-null* sperm (Fig. 4b, left two panels). Moreover, the majority of the top 200 DEGs appeared to be upregulated in the zygotes from *Kit^+/copGFP^* oocytes injected with *Dnmt2-*null sperm. In contrast, the majority of the mRNAs were mostly downregulated in the zygotes from *Kit^+/copGFP^* oocytes injected with WT sperm. These data suggest that zygotes derived WT oocytes are more resistant than those from *Kit^+/copGFP^* oocytes to the impact of sperm genotype (WT *vs*. *Dnmt2*-null).

The *Dnmt2^+/-^* zygotes derived from *Dnmt2*^*-/-*^ sperm and *Kit^+/copGFP^* oocytes and WT zygotes derided from WT sperm and *Kit^+/copGFP^* oocytes mimic the two breeding schemes which showed the totally blocked transmission of the WTT phenotype and the maximal transmission of the WTT phenotype (Fig. 1c and Fig. 4c). Between these two types of zygotes, a total of 1,948 dysregulated genes were identified, including 1,378 upregulated and 570 down-regulated in *Dnmt2^+/-^* zygotes (Fig. 4c and supplemental information, Table S4). GO term enrichment analyses revealed that these dysregulated genes were involved in multiple biological functions, including epigenetic regulation (e.g., histone modifications and chromosome/chromatin organization), energy metabolism, cell signaling and autophagy (Fig. 4d).

To reveal the differences in gene expression between zygotes derived from WT (baseline levels of WTT phenotype) and *Kit^+/copGFP^* oocytes (maximal penetrance of the WTT phenotype) injected with WT sperm, we also compared their transcriptomes and found a total of 1,453 dysregulated genes, including upregulated and down-regulated ones (Fig. 4e and supplemental information, Table S5). Of interests, the GO term enrichment analyses identified that these dysregulated genes, similar to those dysregulated genes between the *Dnmt2^+/-^* zygotes from *Kit^+/copGFP^* oocytes injected with *Dnmt2*-null and WT sperm (Fig. 4d), are Mostly involved in epigenetic regulation, e.g., histone modifications and chromosome/chromatin organization, as well as energy metabolism. More interestingly, among all the DEGs between the two groups, a total of 264 genes were shared, and these shared genes function in multiple cellular processes including histone modifications, signaling and metabolism (Supplemental information, Fig. S7).

### Histone modification genes as the potential targets of the dysregulated tsRNAs and miRNAs in *Dnmt2*-null sperm

Given that the major difference between the two breeding schemes, i.e., a complete block of the WTT phenotype (*Dnmt2*^*-/-*^ male X *Kit^+/copGFP^* female) and the maximal transmission of the WTT phenotype (*Dnmt2+/+* male X *Kit^+/copGFP^* female), lies in the sperm (*Dnmt2*-null vs. WT), the most likely factors responsible for the failure of maternal transmission of the paramutant WTT phenotype would be those DNMT2-regualted sncRNAs, especially tsRNAs. Based on the report that tsRNAs, especially 3’CCA tsRNAs, can act as miRNAs and target mRNA transcripts (*20*), we performed *in silico* sequencing match to identify the potential targets of those differentially expressed, DNMT2-dependent tsRNAs in *Dnmt2*-null sperm (Fig. 5). Following an established method (*21*), we first generated all the possible transcripts isoforms and compared their differential expression between *Dnmt2^+/-^* zygotes from *Kit^+/copGFP^* oocytes injected with *Dnmt2*-null sperm and WT zygotes from *Kit^+/copGFP^* oocytes injected with WT sperm using Stringtie and Cuffdiff. We then used GSTAr (Generic Small RNA-Transcriptome Aligner), a flexible RNAplex-based alignment of sncRNAs (15-26nt) to transcriptome, to predict the potential tsRNA complimentary annealing to mRNAs, as described previously (*22*). We identified 348 genes that could be targeted by differentially expressed, DNMT2-dependent 5’ tsRNAs in *Dnmt2*-null sperm (Supplemental information, Table S7). However, no specific enriched GO terms were found from the 5‘ tsRNAs-targeted genes. In contrast, a total of 386 genes were found to be potentially targeted by those differentially expressed, DNMT2-dependent 3’ tsRNAs in *Dnmt2*-null sperm (Supplemental information, Table S8), and 51 out of 386 target genes were significantly differentially expressed between *Dnmt2^+/-^* zygotes from *Kit^+/copGFP^* oocytes injected with *Dnmt2*-null sperm and WT zygotes from *Kit^+/copGFP^* oocytes injected with WT sperm (Fig. 5a and supplemental information, Table S8). The majority (~73%) of the target genes were up-regulated in *Dnmt2^+/-^* zygotes from *Kit^+/copGFP^* oocytes injected with *Dnmt2*-null sperm, as compared to the WT zygotes from *Kit^+/copGFP^* oocytes injected with WT sperm (Fig. 5a). Consistently, overlapping bar plots showed DNMT2-dependent 3’ tsRNAs could preferentially target transcripts that were up-regulated in the *Dnmt2^+/-^* zygotes from *Kit^+/copGFP^* oocytes injected with *Dnmt2-*null sperm (Fig. 5b). To test the statistical significance of this correlation, the Pearson Chi-Square test was performed (Fig. 5c). The results support a strong, positive correlation between DNMT2-dependent 3’ tsRNAs and up-regulated mRNAs in *Dnmt2^+/-^* zygotes derived from *Kit^+/copGFP^* oocytes injected with *Dnmt2*-nul sperm, as compared to WT embryos from *Kit^+/copGFP^* oocytes injected with WT sperm. GO term enrichment analyses revealed that the potential target genes of *Dnmt2*-dependent 3’tsRNAs are involved in several biological functions, including transcriptional control and histone modifications (Fig. 5d). It is noteworthy that histone modification genes appeared once again to be enriched in the DEGs between *Dnmt2^+/-^* (*Kit^+/copGFP^* oocytes injected with *Dnmt2*-null sperm) and WT (*Kit^+/copGFP^* oocytes injected with WT sperm) zygotes (Fig. 4d), suggesting that sperm-borne, DNMT2-dependent 3’ tsRNAs preferentially target histone modification genes post-transcriptionally.

**Fig. 5.**
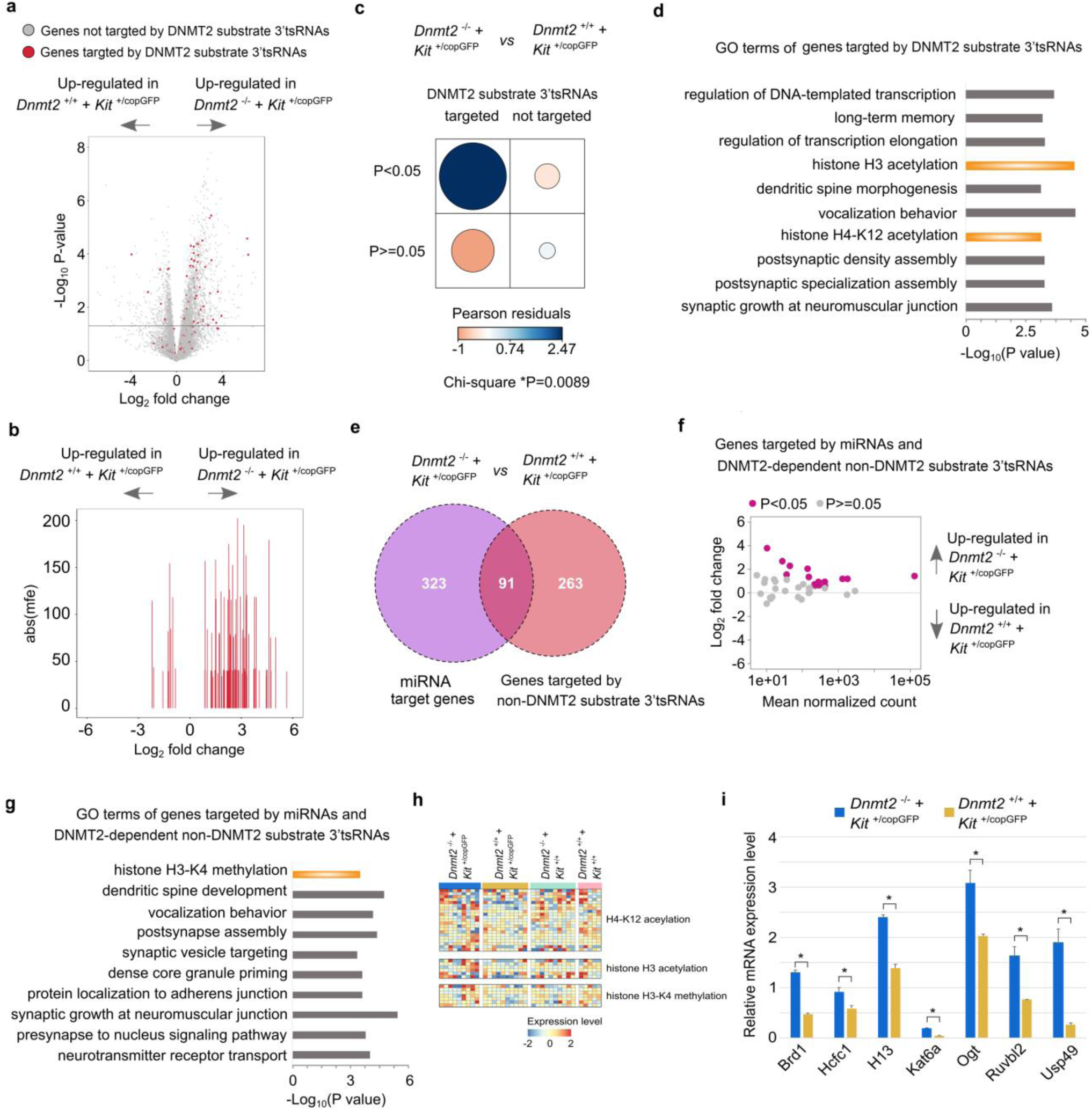
Preferential targeting of genes involved in histone modifications by dysregulated tsRNAs and miRNAs in *Dnmt2*-null sperm. (a) Volcano plots showing differentially expressed genes (DEGs) between zygotes from *Kit*^+/copGFP^ oocytes injected with *Dnmt2*-null (n=9) and WT (n=10) sperm that are also targeted by dysregulated 3’ tsRNAs from DNMT2 substrate tRNAs in *Dnmt2* KO sperm. The *x-axis* shows the Log2 fold change of enriched genes in zygotes from *Kit*^+/copGFP^ oocytes injected with WT or *Dnmt2*-null sperm, whereas the *y-axis* represents the −Log10 P-value. Abline marks the cut-off of p = 0.05. b) Overlapping bar plots showing the relationship between genes enriched in zygotes from *Kit*^+/copGFP^ oocytes injected with WT or *Dnmt2*-null sperm, and the targeting strength of DNMT2-dependent 3’ tsRNAs. The *x-axis* shows the Log2 fold changes of enriched gens in zygotes from *Kit*^+/copGFP^ oocytes injected with WT or *Dnmt2*-null sperm. The *y-axis* represents the absolute binding energy [abs(mfe)] of DNMT2-dependent 3’ tsRNAs. (c) Chi-square test of the association between DNMT2-dependent, 3’tsRNAs-targeted genes and DEGs between zygotes from *Kit*^+/copGFP^ oocytes injected with WT or *Dnmt2*-null sperm (p<0.05, n=10). The dot size is proportional to the absolute value of Pearson residuals indicating the strength of association between 3’ tsRNAs and DEGs. Blue to orange gradients indicate positive to negative association. (d) Gene ontology analyses of DNMT2-dependent 3’tsRNAs-targeted genes. Each target gene is mutually targeted by at least three 3’ tsRNAs. The top 10 enriched gene ontology terms of 3’ tsRNAs-targeted genes are shown in bar graphs with the −Log10 P values indicated. (e) Venn diagram showing the number of genes targeted by DNMT2-regulated, non-DNMT2 substrate 3’ tsRNAs overlap with those targeted by miRNAs. (f) MA plots showing the DEGs between zygotes from *Kit*^+/copGFP^ oocytes injected with *Dnmt2*-null (n=9) and WT (n=10) sperm that are targeted by both DNMT2-regulated, none-DNMT2 substrate 3’ tsRNAs and miRNAs. The *x- axis* shows the mean value of normalized counts between the two groups of zygotes compared. The *y-axis* represents the Log2 of zygotes from *Kit*^+/copGFP^ oocytes and *Dnmt2*-null sperm/zygotes from *Kit*^+/copGFP^ oocytes and WT sperm. (g) GO term enrichment analyses of DEGs between zygotes from *Kit*^+/copGFP^ oocytes injected with *Dnmt2*-null and WT sperm that are targeted by both DNMT2-regulated, none-DNMT2 substrate 3’ tsRNAs and miRNAs. (h) Heat map showing key histone modification genes targeted by DNMT2-dependent 3’ tsRNAs and DNMT2-regulated miRNAs. Rows show the mean-centered and normalized log-expression values of genes grouped by each pathway. For the list of genes analyzed, please see Supplemental information, Table S11. (i) qPCR-based validation of expression levels of seven key histone modification genes, including *Brd1, Hcfc1, H13, Kat6a, Ogt, Ruvbl2* and *Usp49* in zygotes from *Kit*^+/copGFP^ oocytes injected with *Dnmt2*-null (n=9) and WT (n=10) sperm.

Given that other sncRNAs, in addition to tsRNAs, were also dysregulated in *Dnmt2*-null sperm, we further analyzed the potential targeting of mRNAs by miRNAs and non-DNMT2 substrate tsRNAs in *Dnmt2*-null sperm. Among the genes that were predicted to be targeted by miRNAs (323 genes) and non-DNMT2 substrate tsRNAs (263 genes), 91 were shared (Fig. 5e, and supplemental information, Tables S9 and S10). Interestingly, the majority of the 91 genes were significantly up-regulated in *Dnmt2^+/-^* zygotes from *Kit^+/copGFP^* oocytes injected with *Dnmt2*-null sperm, as compared to WT zygotes from *Kit^+/copGFP^* oocytes injected with WT sperm (Fig. 5f). GO term analyses of these 91 genes revealed enriched terms in histone H3-K4 methylation, suggesting that the dysregulated non-DNMT2 substrate tsRNAs and miRNAs may co-regulate histone H3 methylation (Fig. 5g). To visualize the differential expression of genes targeted by DNMT2-regulated small RNAs, we grouped 34 histone modification genes by their GO term functional annotations and compared their expressions in PN3/PN4 stage zygotes (Fig. 5h and supplemental information, Table S11). *Dnmt2^+/-^* PN3/4 zygotes from *Kit^+/copGFP^* oocytes injected with *Dnmt2-*null sperm appeared to display overall higher expression levels compared to WT zygotes from *Kit^+/copGFP^* oocytes injected with WT sperm. In *Dnmt2^+/-^* PN3/4 zygotes from *Kit^+/copGFP^* oocytes injected with *Dnmt2-*null sperm, more than half of H4-K12 acetylation and H3 acetylation genes were up-regulated compared to WT zygotes from *Kit^+/copGFP^* oocytes injected with WT sperm. The upregulation of histone modification genes was also validated using qPCR (Fig. 5i). Together, these data strongly support a correlation between sperm-borne, DNMT2-regulated sncRNAs (e.g., tsRNAs and miRNAs) and numerous mRNAs that encode proteins involved in epigenetic regulations (e.g., histone modifications) during zygotic genome activation in pronuclear embryos.

## DISCUSSION

The phenotypes caused by genetic mutations are relatively homogenous, whereas those caused by epimutations tend to be highly variable (*23, 24*). In our *Kit* paramutation mouse model, the WTT phenotype is highly variable with the WTT length ranging from millimeters to centimeter. Also, even in the pure WT C57Bl6/J breeding colonies, ~30% of the WT mice display the WTT phenotype, but the vast majority are short in length (<3 mm). In the breeding scheme for maternal transmission of the WTT phenotype, *Kit^+/copGFP^* females are mated with pure WT males with black tail tips, and the penetrance of the WTT phenotype among the WT offspring almost double (from ~30% baseline levels to ~59%). More interestingly, the increased penetrance predominantly comes from mice with more severe phenotype (>3 mm of WTT), suggesting that longer WTTs likely represent the paramutant phenotype specifically induced by the *Kit* paramutation on the maternal WT allele. The fact that the increased penetrance of the longer WTT phenotype was abolished in *Kit^+/+^* offspring derived from *Kit^+/copGFP^* females mated with *Dnmt2*-null males strongly suggests that *Dnmt2*-null sperm lack factors required for the transmission of the *Kit* paramutation carried on the WT maternal allele through oocytes. Alternatively, this effect may result from *Dnmt2* heterozygosity because these mice, although WT in *Kit* locus, are heterozygous in *Dnmt2* locus (*Dnmt2^+/-^*). However, it is highly unlikely that *Dnmt2* heterozygosity contributed to the inhibitory effects on the maternal transmission of the WTT paramutation phenotype observed because of the following: 1) Sperm sncRNA profiles are similar in sperm from *Dnmt2^+/-^* and WT males (Supplemental information, Fig. S5). 2) When *Dnmt2^+/-^* males were bred with *Kit^+/copGFP^* females, both *Dnmt2* WT (*Dnmt2+/+)* and heterozygous (*Dnmt2*^*+/-*^) offspring with WT *Kit* displayed a distribution pattern of the WTT phenotype similar to that of offspring from WT male and *Kit^+/copGFP^* female breeding pairs (Supplemental information, Fig. S1). Therefore, the drastic inhibition of transmission of the WTT phenotype must be due to the lack of DNMT2-regulayted sncRNAs in *Dnmt2*-null sperm and the action should take place during the early zygotic stages when zygotic genome activation just gets started. Moreover, the slightly decreased baseline penetrance of the WTT phenotype in offspring (*Dnmt2+/-; Kit+/+*) from *Dnmt2*-null males and WT females is indicative of a suppressive effect on the transmission of the WTT phenotype, which is consistent with notion that *Dnmt2*-null sperm are capable of blocking maternal transmission of the paramutant WTT phenotype in the WT C57Bl6/J colonies. Earlier reports have shown that DNMT2 is required for the paternal transmission of a HFD-induced metabolic disorder and a *Kit* paramutation-induced white tail phenotype fails to be transmitted to offspring when either parent is on *Dnmt2*-null background (*12, 13*). Our finding here strongly suggest that *Dnmt2*-null sperm block the maternal transmission of the *Kit* paramutation-induced WTT phenotype to offspring.

Our sncRNA-BS-seq analyses further support the role of DNMTas a methyltransferase responsible for the 5mC modification on C38 of three major tRNA substrates (*4*). Earlier studies have also shown that *Dnmt2* inactivation results in increased tRNA breakage and consequently enhanced tsRNA production (Jeltsch et al., 2017). However, it remains unclear where exactly the breakage point(s) are, and whether it happens randomly or in a site-specific manner. Our data demonstrate that ablation of DNMT2 causes enhanced breakups of not only the three substrate tRNAs at the position of C38, but also other non-DNMT2 substrate tRNAs around the anticodon loop region (position of 27-45nt). This is an intriguing finding because it suggests that in addition to 5mC at C38, DNMT2 may affect other types of modifications at other positions and consequently, their stability. Indeed, our recent study has shown that decreased m2G in tRNAs represents a secondary effect of *Dnmt2* inactivation in sperm (Zhang et al., 2018). Although a lack of C38 m5C on certain DNMT2 substrates affects translation of a group of proteins in a tRNA-specific manner (*25*), we noticed that while DNMT2 ablation induces global shortening of tsRNAs, the overall levels of total tRNAs appear to be only slightly reduced. Consequently, the overall translation function of tRNAs may not be affected significantly, which may explain why *Dnmt2*-null mice show no obvious phenotype (*6*). Consistent with a previous report (*26*), an inverse relationship between miRNA and tRNA expression profiles in sperm was observed, suggesting that biogenesis of miRNAs and tRNAs/tsRNAs may be interrelated. Indeed, it has been shown that production of certain tsRNAs, just like miRNA biogenesis, are Dicer-dependent (*27*). Thus, competing for Dicer binding and processing may lead to such an inverse relationship in both WT and *Dnmt2*-null sperm. Consistent with published data (*16*), we also observed biased expression of 5’ tsRNAs in both WT and *Dnmt2*-null sperm. This could result from selective degradation of 3’ tsRNAs after tRNAs breakage at C38. Indeed, several studies have shown that although both 5’ tsRNAs and 3’ tsRNAs can co-IP with AGO protein (*28, 29*), 3’CCA tsRNAs tend to be degraded through a unknown mechanism (*20*).

If sperm-borne sncRNAs have an epigenetic role during early embryonic development, they should act prior to or during zygotic genome activation in pronuclear zygotes in mice (*19*). Once zygotic genome is activated, many of the sperm-borne RNAs, including tsRNAs and other sncRNAs, can be, in theory, produced by the zygotic genome. Besides, the massive degradation of both maternal and paternal transcripts after zygotic genome activation, would not allow them to continue to exist and to act for an extended period of time (*30*). Therefore, we chose to use ICSI to produce PN3-PN4 zygotes that mirror the F1 offspring in the four breeding schemes used for studying the penetrance of the paramutant WTT phenotype in live mice. The single zygote RNA-seq allowed us to explore the underlying molecular mechanism through which *Dnmt2*-null sperm block the transmission of maternal transmission of the WTT paramutant phenotype. Hundreds of mRNAs are dysregulated between *Kit^+/+^* zygotes from *Kit^+/copGFP^* oocytes injected with either *Dnmt2*-null or WT sperm. Interestingly, the top 200 most dysregulated genes are mostly upregulated in *Kit^+/+^* zygotes from *Kit^+/copGFP^* oocytes injected with *Dnmt2*-null sperm, whereas these genes displayed a similar expression patterns between *Kit^+/+^* zygotes from WT oocytes injected with either *Dnmt2*-null or WT sperm. These data suggest that the sources of both sperm (*Dnmt2*-null or WT males) and oocytes (from *Kit^+/copGFP^* females) have great influence on the gene expression patterns in PN3-PN4 zygotes. Of great interest, genes involved in histone modifications and chromatin configuration stand out as the potential targets of dysregulated tsRNAs and miRNAs in *Dnmt2*-null sperm, providing a clue on the molecular mechanism underlying the function of the *Dnmt2*-dependent sncRNAs as the potential carrier of epigenetic information.

In summary, our data support DNMT2 as a methyltransferase responsible for the C38 5mC of tRNAs, and a lack of this mark leads to enhanced breakups of not only the three substrate tRNAs at C38, but also the other tRNAs around the anticodon loop region (between positions 27-45nt. Intact function of DNMT2 in the male germline is required for maternal transmission of a Kit paramutation-induced phenotype (i.e., WTT). These sperm-borne, DNMT2-dependent small RNAs appear to regulate genes involved in histone modifications and chromatin organization. Future investigation is warranted to uncover the detailed molecular events and to test the applicability of this rule to other models of epigenetic inheritance.

## METHODS AND MATERIALS

### Animal use and care

All mice used in this study were on C57Bl6/J background and housed in a temperature and humidity controlled, specific pathogen-free facility under a light-dark cycle (12:12 light-dark) with food and water *ad libitum*. Animal use protocol was approved by Institutional Animal Care and Use Committee (IACUC) of the University of Nevada, Reno, and is in accordance with the “Guide for the Care and Use of Experimental Animals” established by National Institutes of Health (1996, revised 2011). The *Kit^+/copGFP^* mice were generated as described (*14, 31*). *Dnmt2* knockout mice were purchased from the Jackson Laboratory (Stock#:006240) and were backcrossed for six generations onto C57Bl6/J background before experiments were carried out.

### Sperm collection

Mature sperm were isolated from the cauda epididymis of male mice as described (*12*). In brief, cauda epididymal sperm were allowed to swim into 1ml HTF medium incubated at 37°C, and an aliquot of 300 µl of swim-up sperm was collected after 1 hour of incubation. Sperm were centrifuged at 100 xg for 10 min and re-suspended in 200 µl of Tris-HCL medium (pH=8.0). The re-suspended sperm was then aliquoted into tubes (50 µl/tube), which were then snap frozen in liquid nitrogen followed by storage in −80°C.

### Intracytoplasmic sperm injection (ICSI) and collection of oocytes and zygotes

Adult (6–8 weeks of age) virgin female WT and *KitcopGFP/+* mice were selected as oocyte donors for ICSI. Superovulation was performed by intraperitoneal injection with 5IU PMSG at 19:30 and intraperitoneal injection with hCG 48 h later. Oocytes were collected 13–14 h after hCG administration. ICSI was performed according to the established protocol (*32*). PN3 or PN4 stage zygotes were collected 8 h after ICSI, as described previously (*19*).

### Small noncoding RNA bisulfite sequencing (sncRNA-BS-seq)

RNAs were isolated from frozen sperm using mirVana miRNA Isolation Kit (Invitrogen). Bisulfite conversion of sncRNAs was conducted using the EZ RNA Methylation Kit (ZYMO Research). Small RNA libraries were constructed using the NEBNext Small RNA Library Prep Set for Illumina (NEB). The sncRNA libraries were sequenced on an Illumina HiSeq4000 sequencer (SE50).

### Bioinformatic analyses of sncRNA-BS-seq data

The sncRNA-BS-seq data have been deposited into the NCBI SRA database (accession#: PRJNA516832). The low-quality sncRNA bisulfite sequencing reads and reads shorter than 15 nt were trimmed by Cutadapt (*33*). Then clean reads were C→T converted. The AASRA pipeline (*17*) with default parameters was used to map the remaining reads to known mouse sncRNA, consisting of mature miRNA (miRBase, release 21), precursor miRNA (miRBase, release 21), tRNA (Genomic tRNA Database), piRNA (piRBase), rRNA (ENSEMBL, release 76), snRNA, snoRNA and mitochondrial RNA (ENSEMBL, release 76). The sncRNA references are also C→T converted prior to alignment. Considering tRNA breakdown, all the reads aligned to tRNA subtype reference sequences are recognized as tRNA reads. The cytosine on tRNA reads are recorded and percentage of m5C are calculated after alignment (methylated count/ all read counts, at each tRNA nucleotide). tRNA alignment positions were tracked, with insertion/deletion and mismatch considered. Then nucleotide position density was plotted to reflect tRNA nucleotide density from 5’ to 3’. To determine small RNA expression levels, the aligned reads were counted by featureCounts (*34*). Read counts outputs from AASRA was normalized and analyzed by DESeq2 (*35*). Similar analyses were conducted using small RNA-seq datasets of oocytes and early embryos (Accession#: SRR15386553 and 1538552) from a previous study (Yang et al., 2016).

### Single zygote genotyping

Since we only wished to study the zygotes with wild-type *Kit*, we genotyped single zygotes based on first observation of copGFP followed by PCR-based detection of copGFP using cDNA libraries constructed for RNA-seq. Specific primers that detect *copGFP* cDNAs were used, as previously described (*31*), and *Gapdh* was used as an internal positive control. Zygotes with *Gapdh* but without *copGFP* expression (*Kit^+/+^*) were chosen for sequencing, whereas those with both *Gapdh* and *copGFP* expression (*Kit^+/copGFP^*) were excluded. For primer sequences, please see Supplemental Table S12.

### Single zygote RNA-seq

The zygotes of PN3-PN4 stages collected were stored in −80°C prior to library construction. The single cell cDNA libraries were made using SMART-Seq v4 Ultra Low Input RNA Kit (Takara). The cDNAs were fragmentized and Illumina Sequencing adaptors were added using Nextera DNA Library Preparation Kit. The single zygote libraries were sequenced on an Illumina NextSeq500 sequencer (PE75).

### Bioinformatic analyses of single zygote mRNA-Seq data

The single zygote RNA-seq data have been deposited into the NCBI SRA database (accession#: PRJNA516834). The raw data was processed by Trimmomatic (v0.33) (*36*) to remove adaptor sequences and filter out the bad quality reads. The clean reads were then aligned to UCSC MM9 mouse genome by HiSAT2 (v2.1.0) (*37*). The raw gene count data were generated by featureCount (*34*) and analyzed by the R package Scater (*38*) to filter out biased samples and get library-size and spike-in normalized gene expression values. Transcripts were assembled by Cufflinks (v2.2.1) (*39*). The enriched genes were sent to gene ontology analysis performed by Gene Ontology Consortium with PANTHER Classification system using default parameters (*40*).

### *In silico* sncRNA target prediction

The RNAplex-based script Generic Small RNA-Transcriptome Aligner from Cleveland 4 package (*41*) was used with default parameter to identify cufflinks-assembled mRNA transcripts targeted by miRNAs and tRNA fragments. The output results were imported into R as a data frame for downstream analyses. The transcripts targeted by at least 3 different miRNAs or tRNA fragments are considered a valid target. The corresponding genes of valid targets were loaded to Gene Ontology Consortium with PANTHER Classification system for gene ontology analysis using default parameters (*40*).

### qPCR analyses of key histone modification genes

qPCR analyses were performed as described (*42*). Primer sequences used can be found in Table S12.

## SUPPLEMENTARY MATERIALS

Supplementary material for this article is available at:

## Acknowledgements

We thank Dr. Qi Chen at University of California, Riverside for critical reading of our manuscripts and for his helpful comments.

## Author contributions

W.Y. designed the research. T.Y. performed sncRNA-BS-seq and single zygote RNA-seq. Y.X. and C.T. conducted bioinformatic analyses; Y.W. performed ICSI and collected all of the PN3/PN4 zygotes; S.Y. and H.Z. conducted all of the breeding and phenotypic analyses; W.Y. wrote the manuscript.

## Data and materials availability

Datasets from single zygote RNA-seq and sncRNA-BS-seq have been deposited into the NCBI SRA database with accession numbers PRJNA516834 and PRJNA516832.

## Supplemental Figures

**Figure S1.**
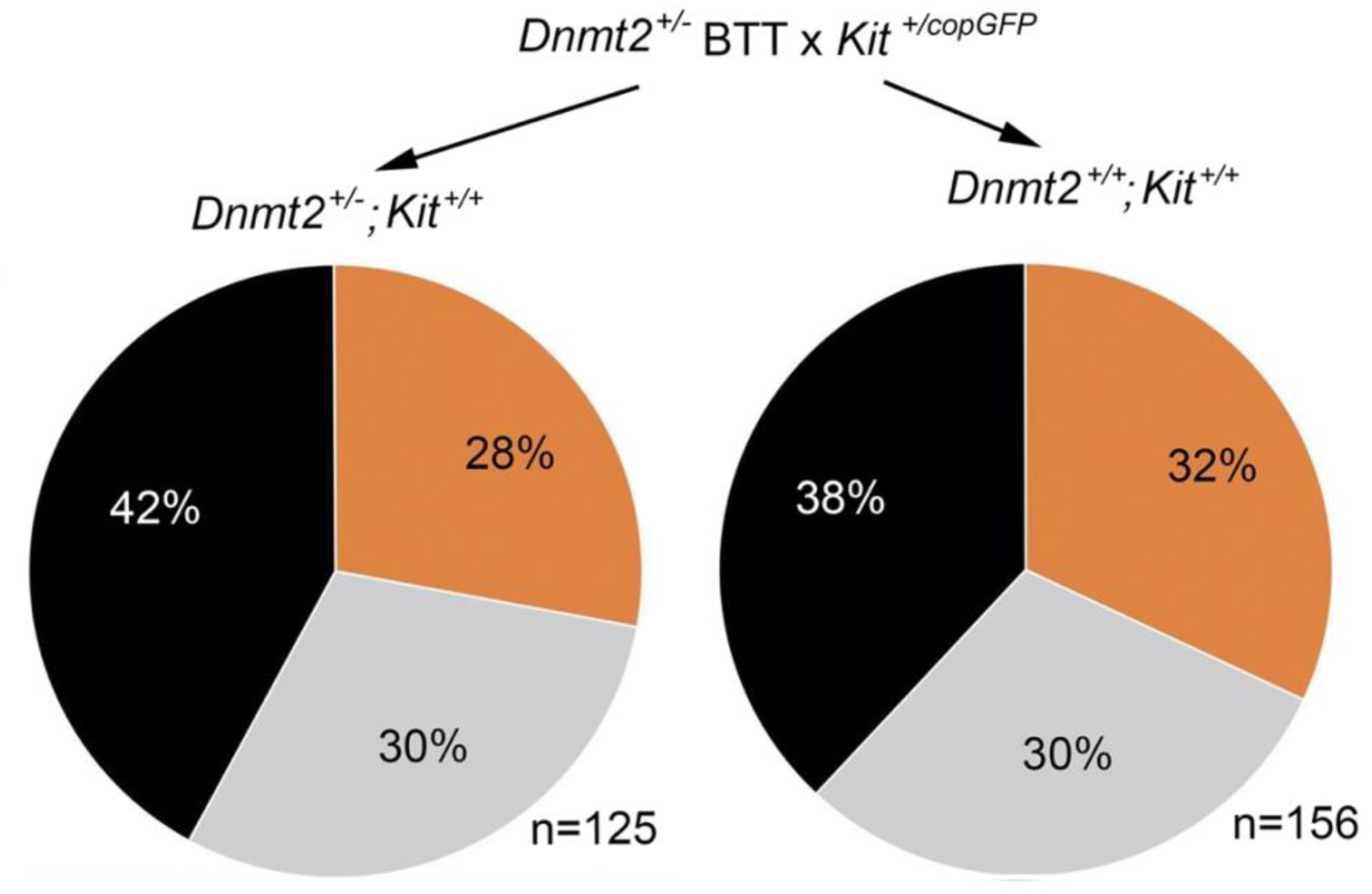
The penetrance of the WTT phenotype in *Dnmt2+/-; Kit+/+ and Dnmt2+/+; Kit+/+* offspring derived from *Dnmt2*^*+/-*^ males and *Kit*^*+/copGFP*^ females.

**Figure S2.**
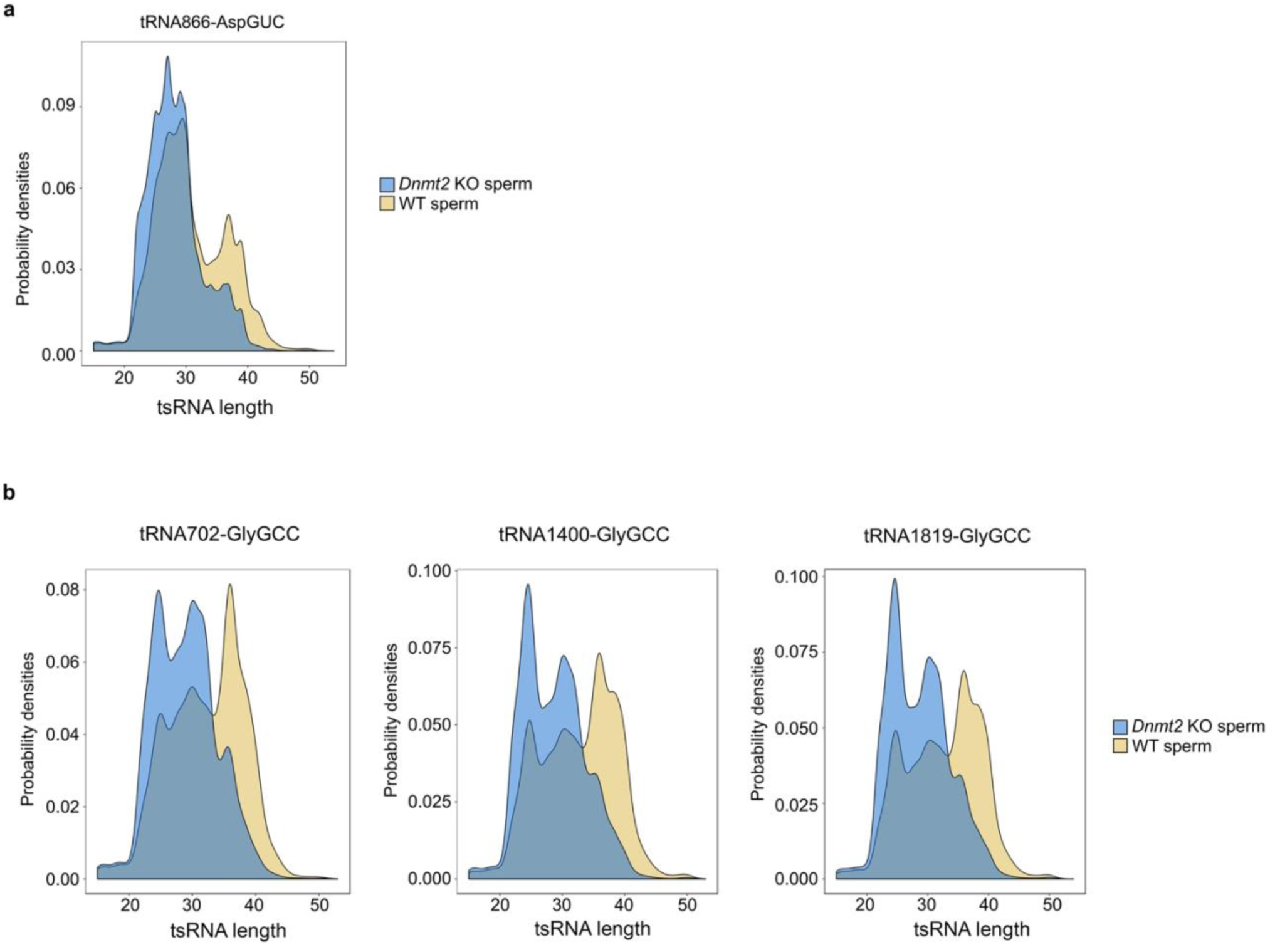
Length distribution of four tsRNAs derived from tRNA-Asp_GUC_ (a) and tRNA-Gly_GCC_ (b) in wild-type (WT) and *Dnmt2* knockout (KO) sperm.

**Figure S3.**
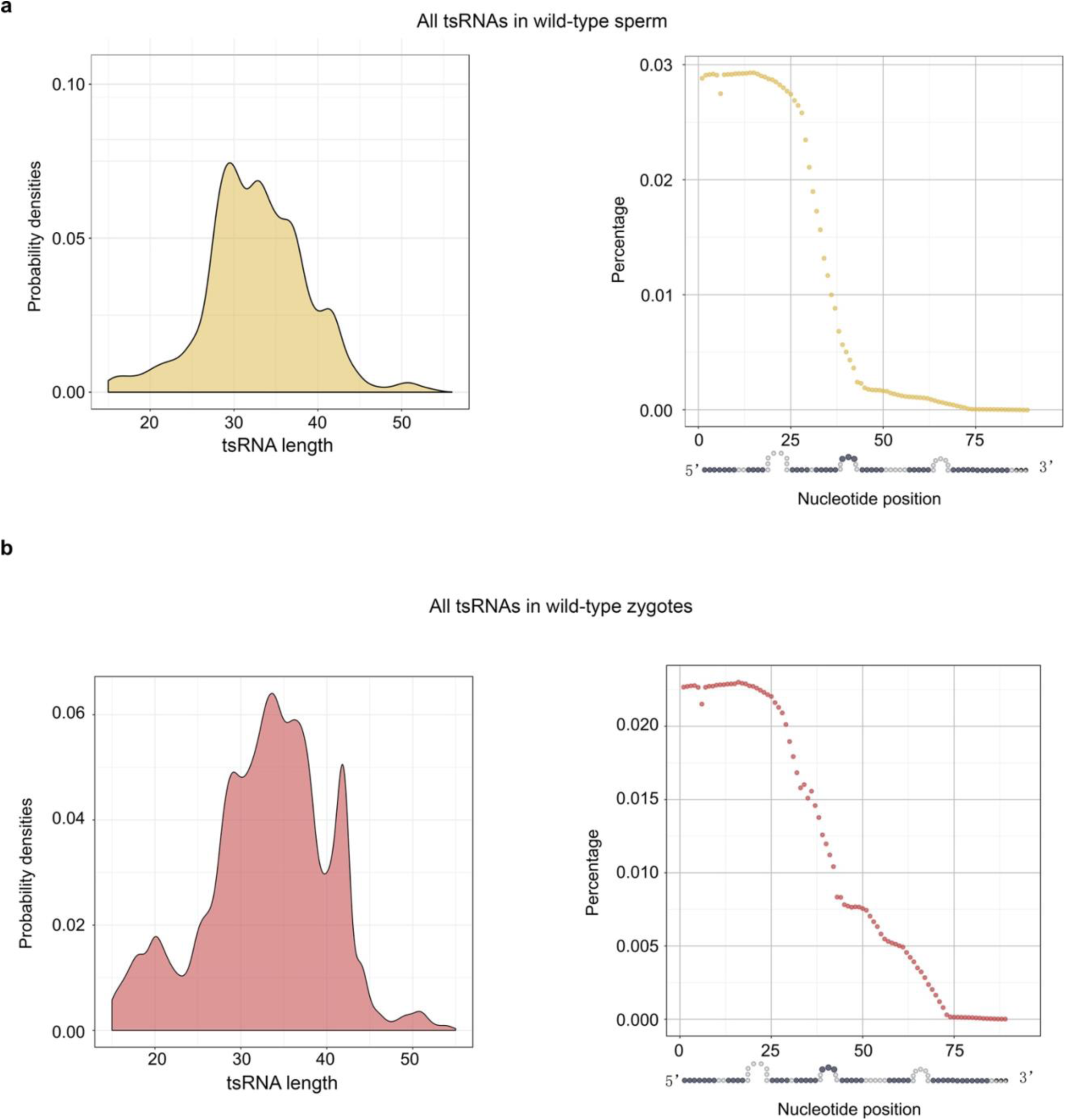
Length distribution and nucleotide position percentage of all tRNAs from wild-type sperm (a) and zygote (b). The datasets used were from a report by Yang, et. al., (Science Advancement, 2016, 2(6); doi: 10.1126/sciadv.1501482).

**Figure S4.**
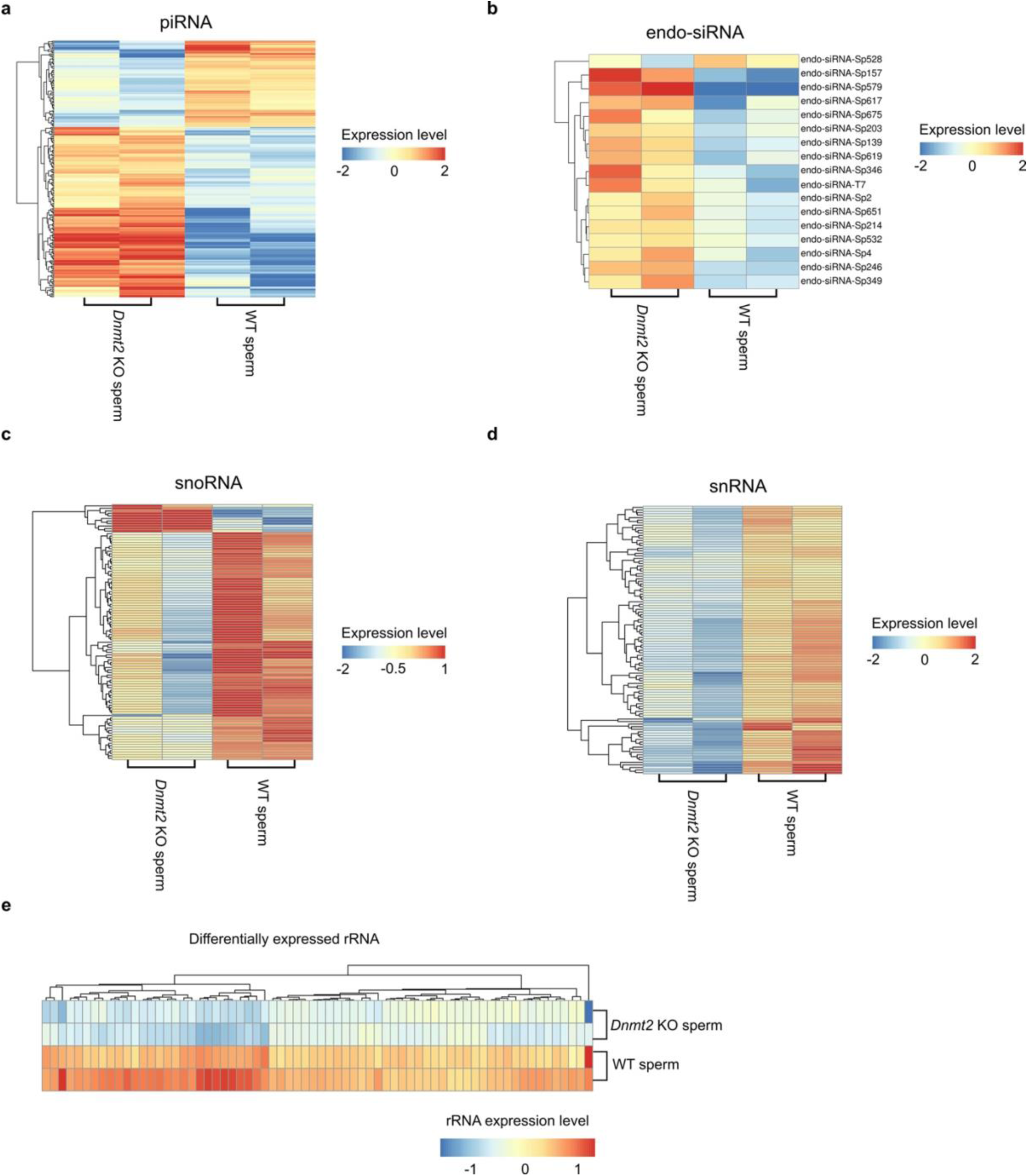
Dysregulated piRNAs (a), endo-siRNAs (b), snoRNAs (c), snRNAs (d), and rRNAs (e) in *Dnmt2* KO sperm compared to WT sperm. For detailed lists of these sncRNAs, please see Supplemental information, Table S2.

**Figure S5.**
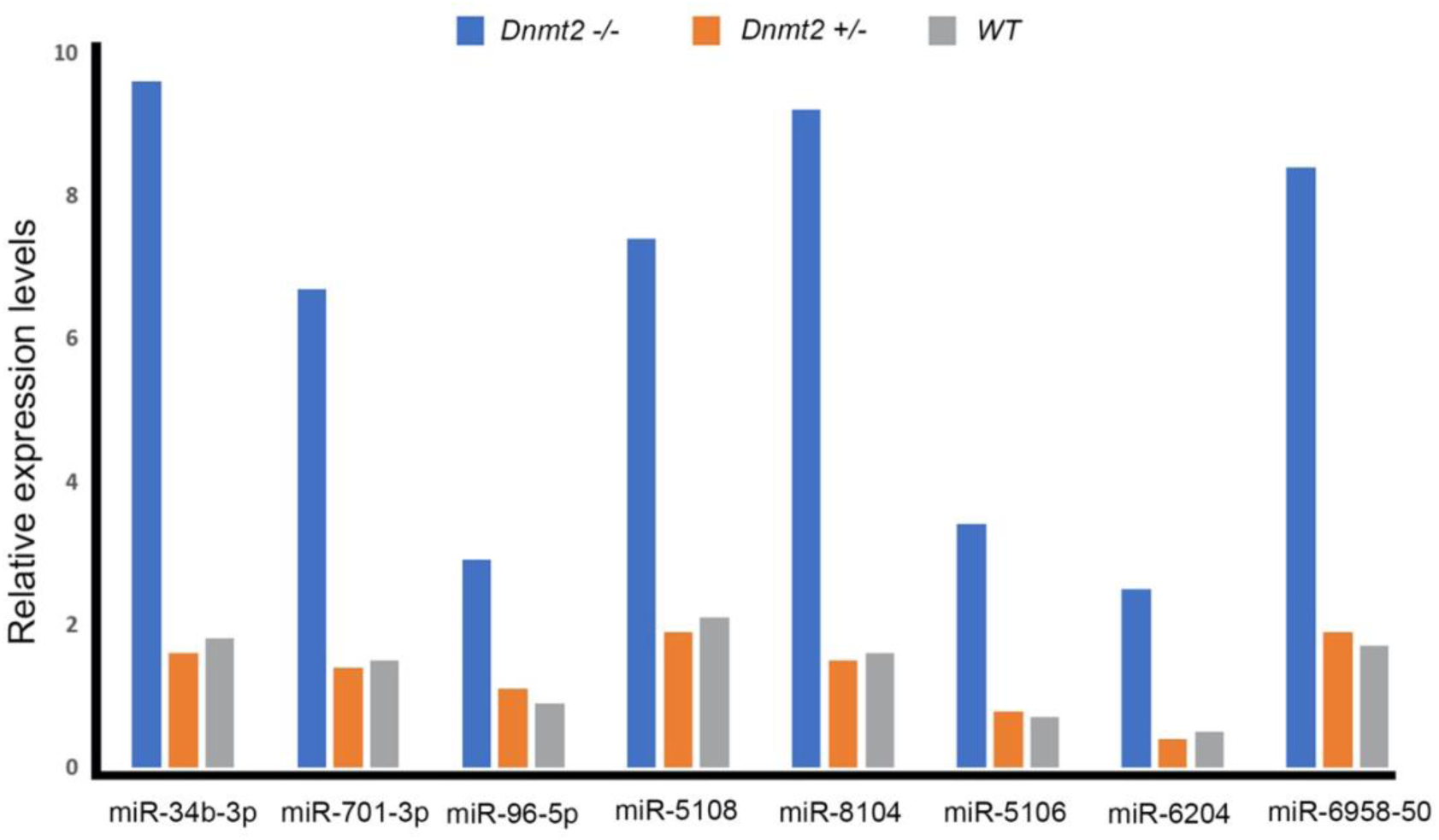
qPCR analyses of the levels of eight miRNAs in wild-type (WT), *Dnmt2^+/-^* and *Dnmt2*^*-/-*^ sperm.

**Figure S6.**
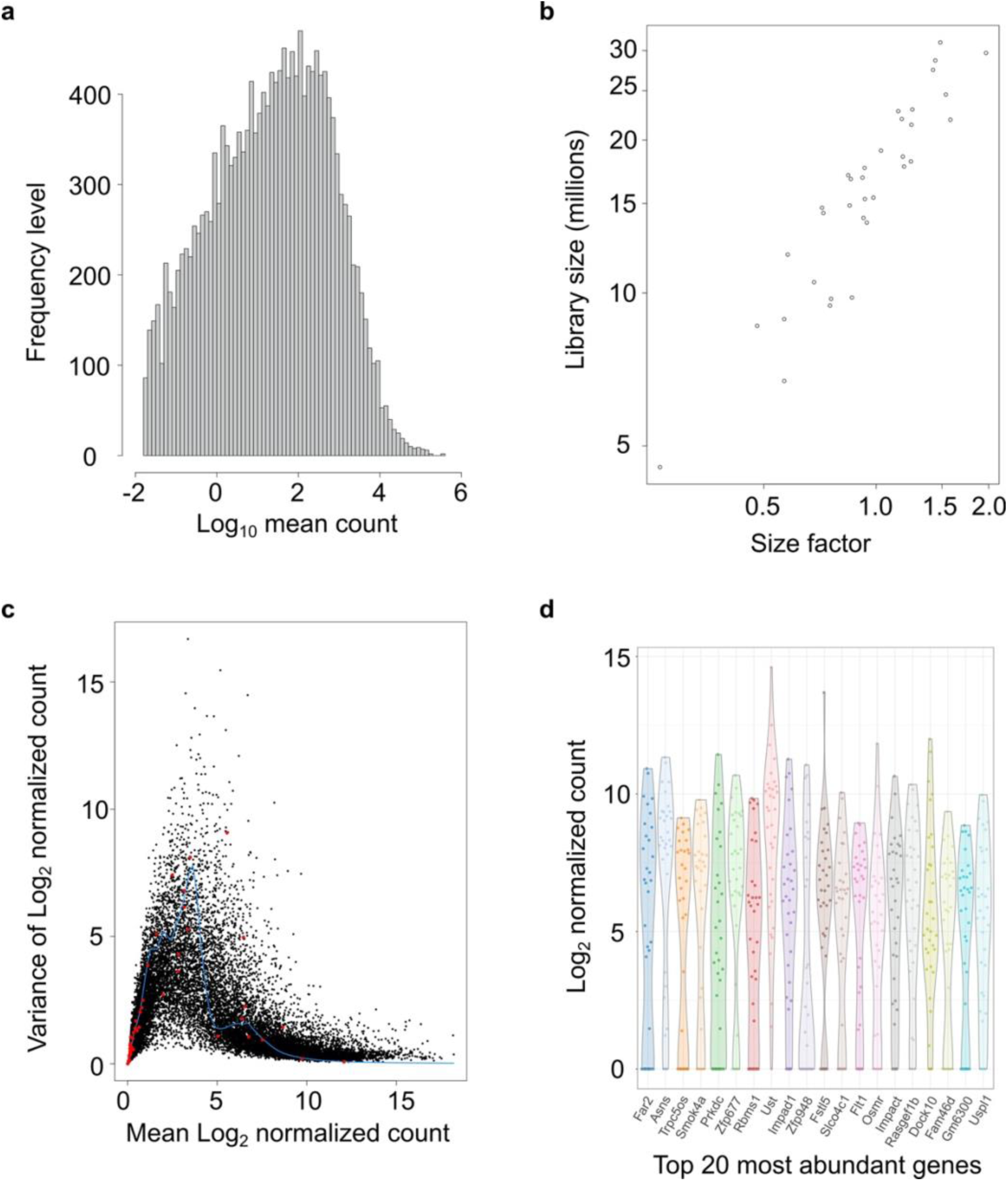
Quality control data of single zygote RNA-seq analyses. (a) Histograms showing the log10 values of mean counts of the filtered gene dataset. (b) Scatter plots with the size factor from deconvolution plotted against the library size for all cells in the dataset. (c) Scatter plots with variance of the filtered and normalized log2 counts plotted against the mean log2 normalized counts. The blue line indicates the mean-dependent trend fitted to the variances of the endogenous gens. The red dots represent the variance estimates for the spike-in transcripts. (d) Violin plots of normalized log-expression for the top 20 highly variable genes in filtered and normalized gene datasets.

**Figure S7.**
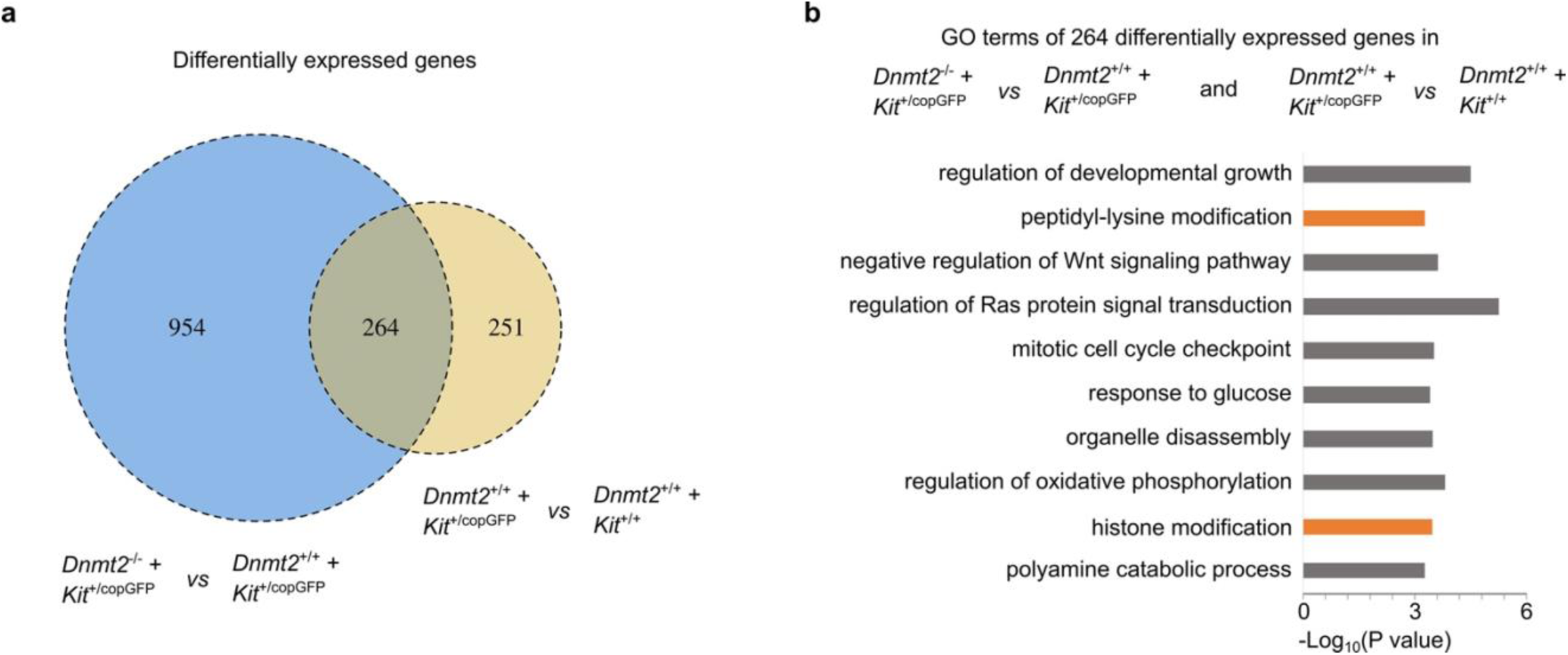
Differentially expressed genes between zygotes derived from *Kit*^+/copGFP^ oocytes injected with *Dnmt2*-null or WT sperm overlap with those between zygotes from *Kit*^+/copGFP^ or WT oocytes injected with WT sperm (a) and GO term enrichment analyses of the 264 overlapping genes (b).

